# Amyloid-Tau PET Fusion Reveals Covarying Network Territories Linked to Disease Stage, Cognition, and APOE4 in Alzheimer’s Disease

**DOI:** 10.64898/2026.08.11.744123

**Authors:** Bikesh Bimali, Jiayu Chen, Armin Iraji

## Abstract

**Background:** Amyloid-beta (Aβ) and Tau are defining pathologies of Alzheimer’s disease, but their relationship varies substantially across the brain. Amyloid deposition is spatially widespread, whereas Tau shows greater regional heterogeneity and closer relationships with disease severity. Consequently, understanding the disease requires more than measuring the two pathologies independently or summarizing them within predefined regions. A major unresolved question is how Amyloid and Tau covary across the whole brain, where their spatial patterns converge or diverge, and whether their local combination carries distinct information about disease stage and cognition.

**Methods:** We analyzed paired florbetapir Amyloid PET and flortaucipir Tau PET from the Alzheimer’s Disease Neuroimaging Initiative (ADNI), comprising 378 paired imaging sessions from 320 participants spanning cognitively normal (CN), mild cognitive impairment (MCI), and Alzheimer’s Dementia (AD). High-order joint independent component analysis was used to identify fine-grained, data-driven patterns of Amyloid-Tau covariance across individuals. The resulting Amyloid and Tau maps were characterized by their spatial similarity and correspondence with rsfMRI-derived intrinsic functional networks. We then used the same data-driven spatial regions to quantify, for each participant, the relative extent of Amyloid abnormality alone, Tau abnormality alone, and spatially overlapping Amyloid-Tau abnormality.

**Results:** Ninety-six non-artifactual Amyloid-Tau components were identified, of which 73 showed appreciable spatial correspondence between their paired Amyloid and Tau maps, while 17 were Tau-localized and 6 Amyloid-localized. Amyloid component maps more frequently corresponded with rsfMRI-derived intrinsic network organization than Tau maps. Expression of the joint components differed across diagnosis groups, demonstrating spatially heterogeneous disease-stage patterns. Within the same data-driven regions, isolated Amyloid, isolated Tau, and overlapping Amyloid-Tau abnormality showed markedly different disease-stage profiles: Amyloid-only abnormality was more prominent in the earlier CN-to-MCI contrast, Tau-only abnormality in contrasts involving AD, whereas spatially overlapping Amyloid-Tau abnormality showed the broadest differences across disease stages and the largest mean CN-to-AD expansion. Overlapping abnormality was also associated with cognition across the greatest number of components and showed the strongest overall association with ADAS13. APOE4-related diagnosis-stage differences were considerably more widespread for pathology measures containing Amyloid than for Tau-only abnormality.

## Introduction

Alzheimer’s disease (AD) is biologically characterized by abnormal Amyloid-beta (Aβ) plaques and Tau neurofibrillary pathology, and contemporary biomarker frameworks increasingly define and stage AD using biological evidence of Amyloid and Tau pathology together with markers of disease severity and neurodegeneration (Jack *et al*., 2018; Jack Jr. *et al*., 2024). Molecular PET imaging enables these proteinopathies to be measured in vivo. Amyloid PET tracers, including florbetapir (FBP), are sensitive to early and spatially widespread Aβ plaque deposition (Clark *et al*., 2011; Landau *et al*., 2012), whereas Tau PET tracers, including flortaucipir (FTP), more closely reflect neurofibrillary tangle distribution, regional disease involvement, and cognitive impairment (Johnson *et al*., 2016; Pontecorvo *et al*., 2019; Ossenkoppele, van der Kant and Hansson, 2022). Paired Amyloid-Tau PET therefore provides complementary information about the disease pathology, but understanding how the two pathologies relate spatially across the brain remains an important challenge.

The Amyloid-Tau relationship is not spatially uniform. Regional imaging studies suggest that local Amyloid burden and brain connectivity are associated with where Tau emerges and subsequently accumulates, while longitudinal Tau PET studies demonstrate substantial variation in the spatial extent and progression of Tau across individuals and disease stages (Lee *et al*., 2022; St-Onge *et al*., 2024; Zheng *et al*., 2024; Fonseca *et al*., 2026). This organization also intersects with large-scale brain systems. Early Amyloid deposition preferentially involves distributed association regions, including core default-mode areas, whereas Tau accumulation has been related to the connectivity and vulnerability of highly connected network regions (Palmqvist *et al*., 2017; Frontzkowski *et al*., 2022). Together, these findings motivate a spatially resolved characterization of coordinated Amyloid-Tau variation rather than treating their relationship as a single global association. A data-driven map of Amyloid-Tau coupling can therefore be examined not only for anatomical correspondence between the paired PET maps, but also for correspondence with intrinsic functional network organization.

Most PET studies characterize Amyloid and Tau using global summary measures, atlas-defined regions of interest, Braak-based composites, longitudinal regression models, or connectivity-informed models of Tau spread (Cho *et al*., 2016; Franzmeier *et al*., 2020; Vogel *et al*., 2020; St-Onge *et al*., 2024). These approaches have established important relationships among Amyloid, Tau, network architecture, neurodegeneration, and cognition. However, they generally begin with predefined anatomical regions, network labels, or propagation models. As a result, they are less suited to discovering latent whole-brain spatial patterns in which Amyloid and Tau covary across individuals without imposing the spatial organization beforehand. Moreover, identifying cross-subject covariance alone does not establish whether the paired Amyloid and Tau expressions of that covariance occupy the same anatomical territory or remain spatially distinct. Distinguishing these properties is important for characterizing the spatial organization of coupled pathology.

Multimodal fusion provides a data-driven framework for addressing this problem. Joint independent component analysis (joint ICA) identifies linked sources of variation across modalities through shared subject-level component expression while retaining modality-specific spatial maps (Calhoun *et al*., 2006; Sui *et al*., 2012). The framework continues to be applied in contemporary multimodal neuroimaging, including fusion of Amyloid PET with functional network connectivity in AD (Bimali *et al*., 2025) and joint-ICA-based integration of genomic and functional-connectivity features (Chen *et al*., 2026). One reason joint ICA is particularly well suited to paired Amyloid-Tau PET is that it links the two modalities through shared cross-subject component expression without constraining their modality-specific maps to occupy the same anatomical locations. Amyloid and Tau can therefore covary across individuals as part of the same joint component while showing either spatially corresponding or distinct regional patterns, allowing their spatial relationship to emerge from the data rather than being specified a priori.

Prior Amyloid-Tau PET fusion work established the feasibility of this approach. (Pereira *et al*., 2019) applied joint component analysis to Amyloid and Tau PET and identified broad patterns in which the two tracers showed both related and distinct spatial organization. However, fine-grained whole-brain mapping of paired Amyloid-Tau covariance remains underexplored. ICA model order influences the spatial scale at which these patterns are represented: lower-order solutions capture broader sources, whereas higher-order decompositions can subdivide these sources into more spatially specific components (Abou-Elseoud *et al*., 2010; Iraji *et al*., 2023).

The present study used this framework to address two connected aspects of paired Amyloid-Tau pathology. First, we generated a fine-grained, data-driven map of cross-subject Amyloid-Tau covariance and characterized the spatial organization of each joint component by determining whether its paired Amyloid and Tau maps were spatially co-localized or modality-localized and by relating these maps to rsfMRI-derived intrinsic functional network organization. We hypothesized that, because Amyloid and Tau are biologically related in AD, many covarying Amyloid-Tau components would localize to the same or closely related territories, even though joint ICA does not impose spatial overlap between modalities. At the same time, we expected the high-order decomposition to reveal modality-localized components reflecting distinct spatial behavior of Amyloid and Tau.

Second, we used the spatial territories discovered by joint ICA as common units for subject-specific pathology characterization. This step addresses a complementary question to component covariance: once a territory of coordinated Amyloid-Tau variation has been identified at the population level, how is that territory occupied by Amyloid and Tau abnormality within an individual? We therefore quantified Amyloid-only, Tau-only, and overlapping Amyloid-Tau (dual-pathology) abnormality within each component territory and examined their relationships with diagnosis stage, cognition, and APOE4 carrier status. By linking unsupervised multimodal spatial discovery with subject-specific pathology composition within the same data-derived territories, the study was designed to characterize both where Amyloid and Tau covary and how pathology is spatially expressed within those covarying territories across AD.

## Materials and Methods

### Study overview and terminology

The analysis was organized into three connected stages (Figure 1). First, paired Amyloid and Tau PET were decomposed using joint ICA to discover multimodal components defined by coordinated variation across paired imaging sessions. Second, the paired Amyloid and Tau spatial maps of each joint component were characterized by their within-component spatial similarity. Third, the spatial support of each joint component was used as a data-driven territory in which subject-specific Amyloid-only, Tau-only, and dual-pathology burden was quantified and related to diagnosis stage, cognition, and APOE4 status.

**Figure 1.**
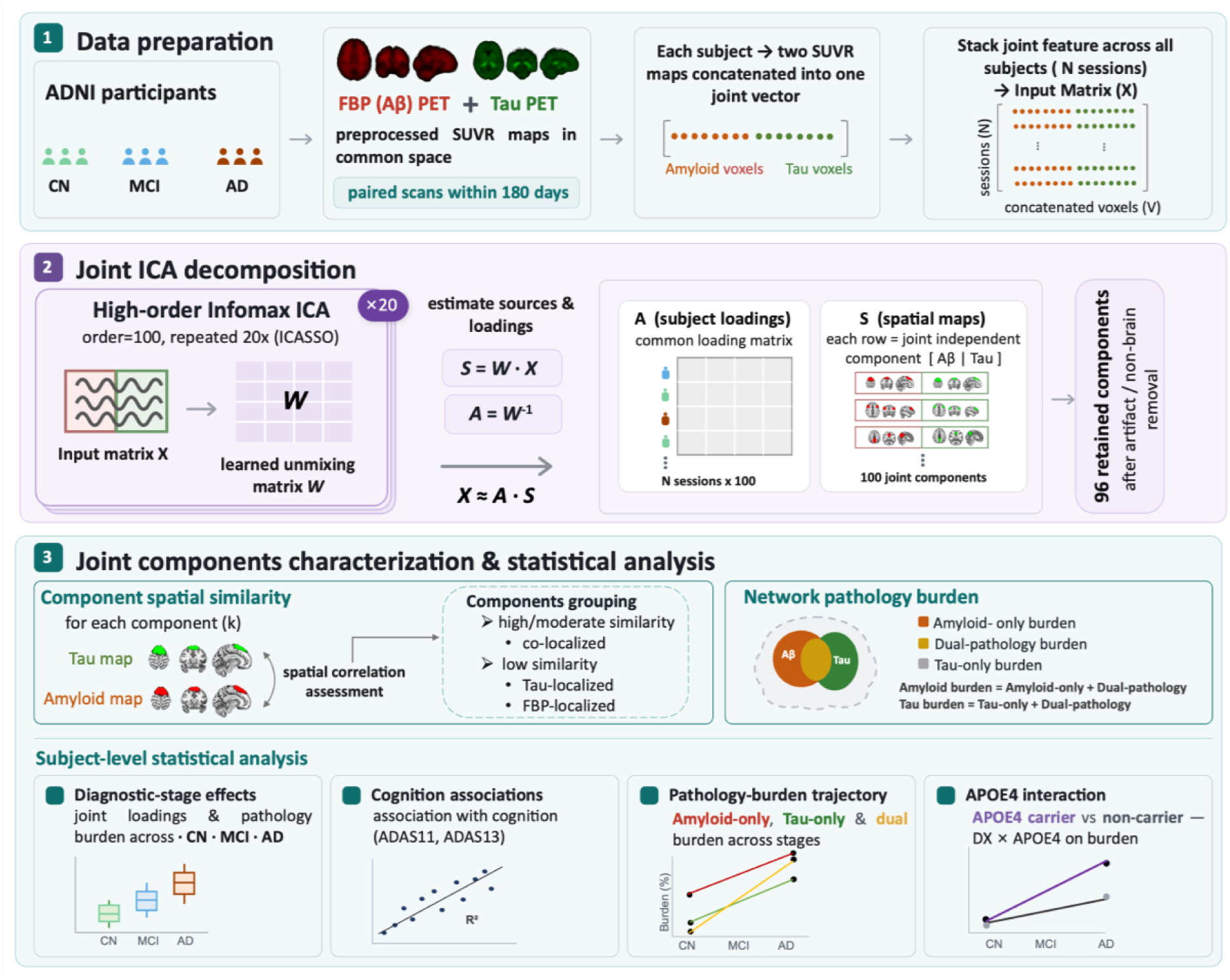
Conceptual overview of the analysis. Paired Amyloid and Tau PET from each imaging session are concatenated and decomposed using joint ICA, yielding a shared session-level loading for each joint component and paired modality-specific spatial maps. The positive spatial support of each joint component defines a component territory, within which subject-specific voxels are partitioned into mutually exclusive Amyloid-only, Tau-only, and dual-pathology burden states. CN, cognitively normal; MCI, mild cognitive impairment; AD, Alzheimer’s disease dementia; ICA, independent component analysis.

Throughout the manuscript, a joint component refers to one ICA-derived multimodal source with a shared session-level loading and paired Amyloid and Tau spatial maps. Component localization: co-localized, Amyloid-localized, or Tau-localized, describes the spatial relationship between these paired ICA maps. For each joint component, a single component territory is then defined from the combined positive suprathreshold spatial support of its Amyloid and Tau maps. Within each territory, Amyloid-only, Tau-only, and dual-pathology burden denote mutually exclusive proportions of voxels showing subject-specific abnormality. Thus, localization terms describe the relationship between the paired ICA maps, the component territory defines one common spatial region for each component, and burden terms describe pathology composition within that region.

### Participants and image preprocessing

Data were obtained from the Alzheimer’s Disease Neuroimaging Initiative (ADNI; adni.loni.usc.edu). Participants were included when both florbetapir Amyloid PET and flortaucipir Tau PET were available within 180 days. The final dataset included 378 paired PET sessions from 320 unique participants spanning CN, MCI, and AD groups (Table 1), with a mean Amyloid-Tau scan interval of 23.01 ± 29.60 days. Most participants contributed a single paired session; 48 contributed two sessions and 5 contributed three sessions. Consequently, the dataset contained limited repeated longitudinal observations, which were not sufficient for modeling within-person longitudinal trajectories.

**Table 1.** Participant characteristics by diagnosis group.

| Diagnosis | N (N*) | Age,<br>mean ± SD | Education,<br>mean ± SD | Female/<br>Male | APOE4<br>negative | APOE4<br>positive | APOE<br>unavailable |
| --- | --- | --- | --- | --- | --- | --- | --- |
| CN | 215 (177) | 72.63 ± 7.53 | 16.71 ± 2.31 | 105 / 72 | 96 | 56 | 25 |
| MCI | 110 (96) | 75.44 ± 7.44 | 15.98 ± 2.83 | 41 / 55 | 46 | 35 | 15 |
| AD | 53 (47) | 76.98 ± 8.20 | 15.47 ± 2.40 | 20 / 27 | 23 | 21 | 3 |

*N\** = unique participants; *N* = paired PET sessions/datapoints used in the imaging analysis; CN = cognitively normal; MCI = mild cognitive impairment; AD = Alzheimer's dementia. APOE4 positive indicates at least one APOE4 allele; APOE4 negative indicates zero APOE4 alleles. Demographic values are based on unique participants (*N\**).

PET images obtained for this analysis were spatially normalized to MNI305 space, expressed as standardized uptake value ratio (SUVR) maps, smoothed to a common 10-mm full-width-at-half-maximum resolution, and resampled to 3-mm isotropic voxels before multimodal analysis. A common whole-brain mask was applied to both modalities, yielding 68,235 voxels per modality. ADNI PET processing and multisite harmonization procedures are described in (Landau *et al*., 2024)

### Joint Multimodal Matrix Construction

For each paired PET session (i), let **f***_i_* ∈ ℝ^1*xV*^ denote the Amyloid voxel vector and ***τ****_i_* ∈ ℝ^1*xV*^ denote the Tau voxel vector, where *V* = 68,235 voxels. The joint multimodal vector for session (i) was defined by concatenating the two modality-specific voxel vectors:

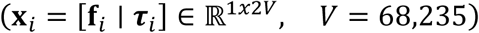

Stacking all paired PET sessions produced the multimodal data matrix (X):

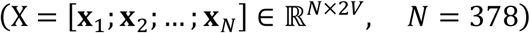

Thus, the input matrix contained 378 paired imaging sessions and 136,470 PET features per session, with the first 68,235 features corresponding to Amyloid SUVR values and the second 68,235 features corresponding to Tau SUVR values. This joint SUVR matrix was decomposed using joint ICA.

### High-Order Joint ICA

Joint ICA modeled the concatenated multimodal matrix as:

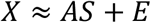

where *A* ∈ ℝ*^N^*^×*K*^contains the session-level joint loading coefficients, *S* ∈ ℝ*^K^*^×2*V*^contains the K multimodal spatial sources, and *E* represents residual error. The decomposition identifies joint sources of coordinated cross-session variation in Amyloid and Tau PET while maximizing statistical independence among the estimated multimodal sources. Because the same loading matrix is shared across the two modality blocks, each component captures an Amyloid-Tau pattern whose expression varies jointly across paired PET sessions.

For component k, the k-th row of S can be separated into paired modality-specific maps 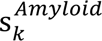 and 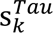, while column k of A provided the shared loading across sessions. Thus, the joint loading represents how strongly the multimodal covariance pattern is expressed in each paired session, whereas the two modality-specific maps indicate where Amyloid and Tau contribute spatially to that pattern. This formulation allows paired Amyloid and Tau maps within the same joint component to show either overlapping or distinct spatial organization.

A model order of K = 100 was selected to characterize Amyloid**-**Tau covariance at a fine spatial scale rather than restricting the decomposition to a small number of broad covariance patterns. Higher model orders can partition broad ICA sources into more spatially specific components (Abou-Elseoud *et al*., 2010), although model-order behavior is dataset and modality-dependent. ICASSO was performed using 20 ICA repetitions to evaluate the reproducibility of the estimated components across repeated decompositions (Himberg and Hyvarinen, 2003). Components whose spatial maps were dominated by CSF or non-brain spatial structure were identified as artifactual and excluded from subsequent analyses, leaving 96 retained joint components.

### Component Spatial Characterization

For each retained joint component, spatial correspondence between the paired Amyloid and Tau maps was quantified using Pearson spatial correlation across the common voxel space. Joint ICA produces continuous-valued spatial maps, in which each voxel reflects the relative contribution of that location to the corresponding component rather than membership in a binary region. Pearson correlation was therefore used to characterize the similarity of the continuous spatial patterns of the paired Amyloid and Tau maps:

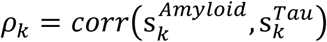

where *ρ_k_* represents the spatial map similarity between the Amyloid and Tau maps of component *k*. Inspection of the empirical distribution of *ρ_k_*values suggested a multimodal structure with approximately three similarity modes. A three-component Gaussian mixture model was therefore fitted to characterize these low, intermediate, and high-similarity regimes. The fitted distributions had mean correlations of 0.278, 0.555, and 0.710, respectively. The crossover between the low- and intermediate-similarity distributions occurred at *ρ_k_* = 0.401, whereas the intermediate-to-high crossover occurred at *ρ_k_* = 0.630. Because the downstream analysis primarily required distinguishing components with clearly low Amyloid-Tau correspondence from those showing appreciable spatial similarity, the first crossover *ρ_k_* > 0.4 was used as the operational threshold. The second crossover characterized variation within the higher-similarity range but was not used to define an additional component class.

This threshold was then used to group the joint components according to the spatial relationship between their paired modality maps. Components with *ρ_k_* > 0.4 were classified as co-localized, indicating appreciable spatial correspondence between the Amyloid and Tau maps. Components with *ρ_k_* ≤ 0.4 were classified as modality-localized and further designated as Amyloid-localized or Tau-localized according to which modality showed the clearer focal spatial pattern. These terms describe the relative spatial organization of the paired maps and do not imply absence of signal in the other modality.

### Alignment with rsfMRI-derived intrinsic functional networks

After grouping components according to their Amyloid-Tau spatial relationship, we next assigned each component an interpretable functional-system label. Because the joint ICA components were identified without predefined anatomical or functional labels, the modality-specific Amyloid and Tau maps were compared with NeuroMark 2.2 intrinsic connectivity network templates derived from large-scale resting-state fMRI data (Iraji *et al*., 2023; Jensen *et al*., 2024). NeuroMark provides continuous spatial ICN maps, allowing direct spatial comparison with the continuous component maps obtained from ICA.

For each joint component, the Amyloid and Tau maps were correlated separately with the NeuroMark templates, and the closest-matching ICN subdomain was identified. For co-localized components, the shared matching functional domain of the paired maps was used as the component label. For Amyloid-localized or Tau-localized components, the label was assigned according to the NeuroMark match of the modality map that showed the localized spatial pattern. This labeling was intended to provide a recognizable functional context for otherwise data-driven PET components rather than to assume that the PET components themselves represent functional networks. Comparing the two modality-specific maps separately also allowed us to determine whether the Amyloid and Tau expressions of the same joint component were most closely associated with the same or different intrinsic functional systems.

### Component-level pathology composition/burden

To translate population-level joint components into subject-specific measures of regional pathology, we next quantified Amyloid and Tau abnormality within each retained joint component territory. First, for each component *k*, the Amyloid and Tau spatial maps were z-scored, and the component territory *T_k_* was defined as the union of suprathreshold positive spatial support (*z* > 1.96; corresponding to the two-tailed significance value of *p* < 0.05) from either modality:

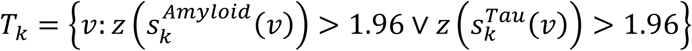

Subject-specific Amyloid and Tau abnormalities were then quantified separately using modality-specific voxelwise thresholds derived from cognitively normal, Amyloid-negative (CN Aβ-) participants. This group was used as a common low-AD-pathology reference. For each voxel v, the Amyloid and Tau abnormality thresholds denoted by *θ* were defined as the 95th percentile of the distribution in the reference group.

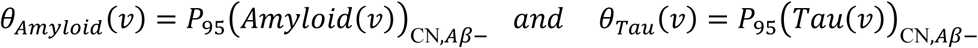

For session *i*, component *k*, and voxel *v* ∈ *T_k_*, abnormality indicators were defined as:

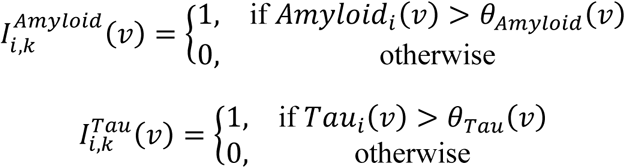

where the indicator equals 1 if the voxel is abnormal for that modality and 0 otherwise.

Three mutually exclusive pathology burden features were then calculated as percentages of the component territory. Amyloid-only burden quantifies the percentage of the component territory where Amyloid is abnormal, but Tau is not abnormal:

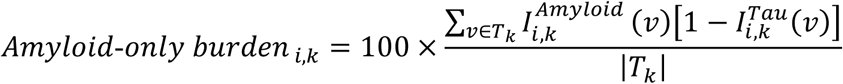

Tau-only burden quantifies the percentage of the territory where Tau is abnormal, but Amyloid is not abnormal:

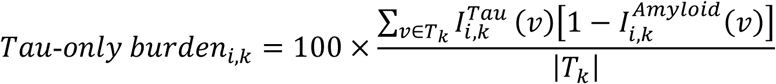

Dual pathology burden quantifies the percentage of the territory where both Amyloid and Tau are abnormal in the same voxels:

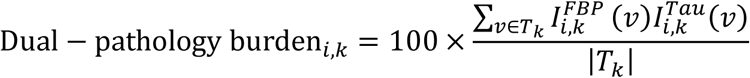

All three measures use the same denominator and therefore represent the subject-specific proportion of a data-driven component territory occupied by a particular pathology state. Amyloid-only and Tau-only burden explicitly exclude voxels in which both tracers are abnormal; those voxels contribute to dual-pathology burden.

### Statistical analysis

All analyses were performed across the 96 retained components. We first tested diagnosis-stage differences in joint component loadings using linear mixed-effects models with diagnosis as the predictor of interest, age, sex, education, race, and imaging site as fixed-effect covariates, and participant identifier as a random intercept to account for repeated sessions:

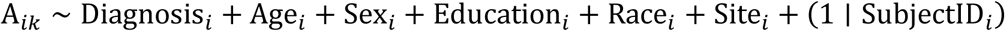

where *i* indexes the paired PET session and *k* indexes the component. Pairwise diagnosis contrasts were evaluated for MCI versus CN, AD versus MCI, and AD versus CN. False discovery rate (FDR) correction was applied across joint components separately within each diagnosis contrast(Benjamini and Hochberg, 1995).

We next examined diagnosis-stage differences in the pathology composition of the discovered component territories. For each component, Amyloid-only, Tau-only, and dual-pathology burden were analyzed using the same linear mixed-effects framework as the joint loadings:

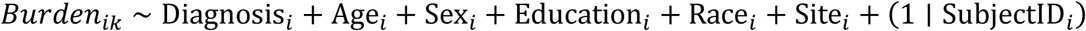

Cognition associations were tested using nested covariate-adjusted models for available cognitive outcomes, including ADAS13, ADAS11, and CDRSB. For each cognitive outcome *y_i_* and component-level predictor *x_ik_*, the base and feature-added models were:

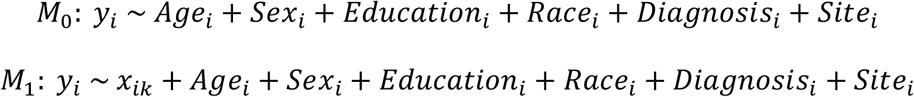

Partial *R*^2^ was calculated from the difference in model *R*^2^ between the feature-added model and the base covariate model. This quantified the variance explained by each component feature beyond demographic, diagnosis, and site covariates. Predictors *x_ik_* included joint loading, Amyloid-only burden, Tau-only burden, and dual-pathology burden.

Finally, APOE4 carrier status was evaluated as a modifier of pathology burden across disease stage. APOE4 carrier status was coded as positive for participants with at least one APOE4 allele and negative for participants with zero APOE4 alleles. Participants with unavailable APOE genotype were excluded from this analysis. For each component and burden feature, the overall diagnosis-by-APOE4 interaction was tested while adjusting for age, sex, education, race, and imaging site:

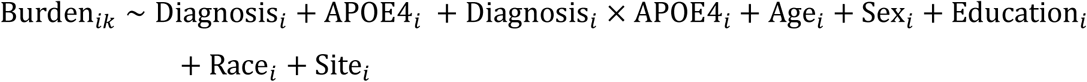

The overall interaction p-value was obtained and FDR-corrected across the 96 components within each burden feature.

## Results

### High-order joint ICA resolves fine-grained Amyloid-Tau coupling territories across intrinsic brain systems

Joint ICA estimated 100 joint components. ICASSO assessment across repeated decompositions yielded a mean component quality index (*I_q_*) of 0.741 ± 0.208, with a median of 0.798 and a range of 0.304-0.986. The ICA decomposition accounted for 94.3% of the variance in the original Amyloid SUVR data and 95.2% of the variance in the original Tau SUVR data, indicating comparable representation of the two PET modalities. Four components dominated by CSF or non-brain spatial structure were identified as artifactual and excluded, leaving 96 components for subsequent analyses. The retained components showed regionally organized covariance patterns across a broad range of brain territories (Figure 2).

**Figure 2.**
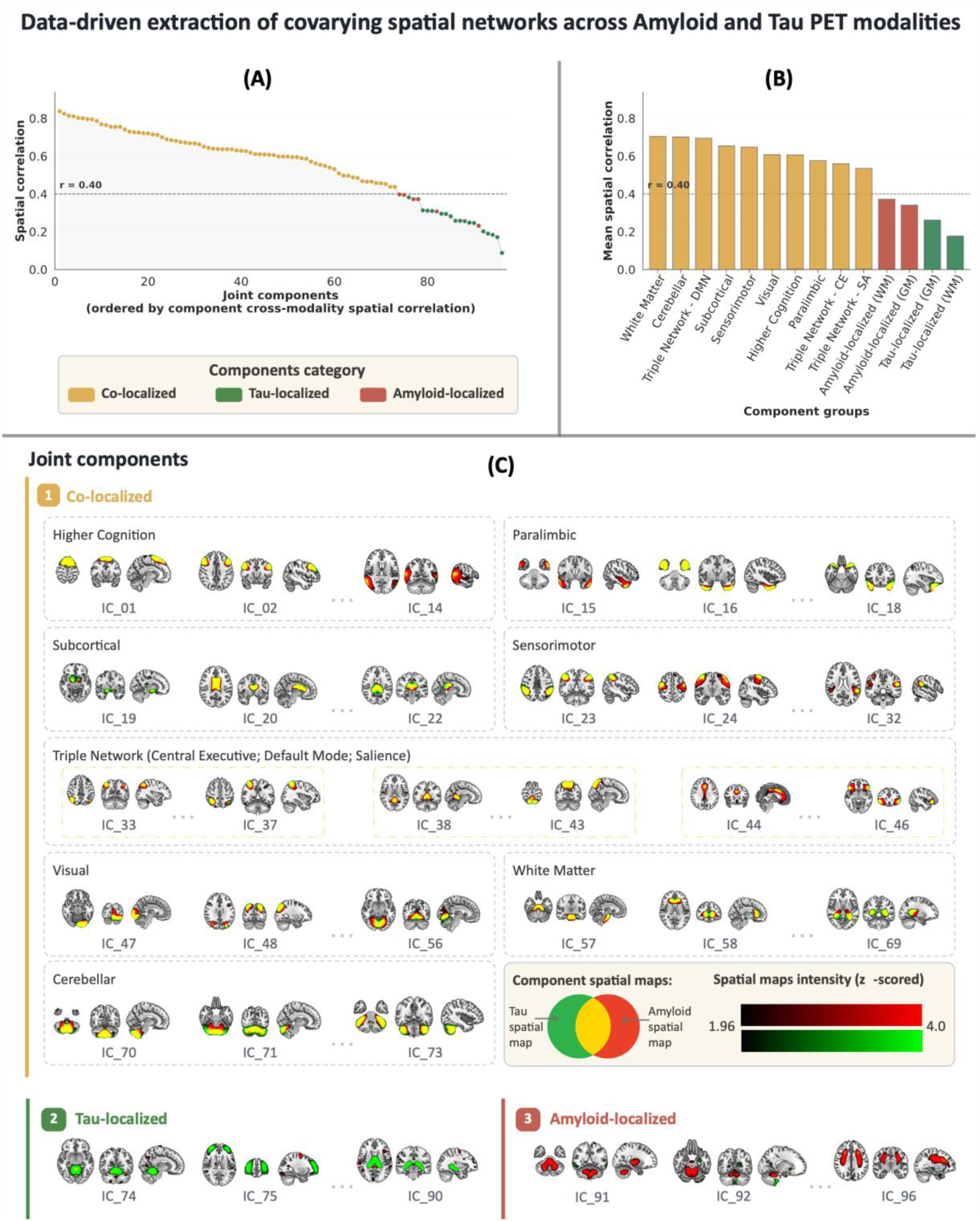
Spatial characterization of the retained joint Amyloid-Tau PET components. (A) Components ordered by Pearson spatial correlation between paired Amyloid and Tau maps. The dashed line at r = 0.40 indicates the data-driven boundary used to identify components with at least moderate Amyloid-Tau spatial similarity. (B) Mean Amyloid-Tau spatial correlation summarized by component grouping. (C) Representative spatial maps illustrating co-localized, Tau-localized, and Amyloid-localized components.

Spatial similarity between the paired Amyloid and Tau maps varied substantially across components (mean ± SD = 0.551±0.185 ; median = 0.598; range = 0.087–0.838). Using the data-driven grouping procedure described in Methods, 73 components were classified as co-localized, whereas 23 showed lower Amyloid-Tau spatial similarity and were classified as modality-localized, including 17 Tau-localized and 6 Amyloid-localized components (Figure 2A-C).

When the component maps were compared with NeuroMark 2.2 rsfMRI-derived intrinsic connectivity network templates, the 73 co-localized components spanned a broad range of NeuroMark-defined systems. In gray matter, co-localized components showed strong alignment with established intrinsic functional networks, including higher-cognition frontal regions (HC-FR; IC1-IC10), higher-cognition insular-temporal regions (HC-IT; IC11-IC13), higher-cognition temporoparietal cortex (HC-TP; IC14), paralimbic regions (PL; IC15-IC18), subcortical basal ganglia and extended thalamic regions (SC-BG/SC-ET; IC19-IC22), sensorimotor cortex (SM; IC23-IC32), triple-network central-executive, default-mode, and salience regions (TN-CE/TN-DMN/TN-SA; IC33-IC46), visual occipital and occipitotemporal regions (VI-OC/VI-OT; IC47-IC56), and cerebellar regions (CB; IC70-IC73). White-matter components included cerebellar-brainstem, frontal, insular, occipitotemporal, sensorimotor, and temporoparietal territories (IC57-IC69).

The 17 Tau-localized components (IC74-IC90) spanned cerebellar, frontal, temporoparietal, paralimbic, subcortical, sensorimotor, triple-network, visual, and white-matter territories, whereas the six Amyloid-localized components (IC91-IC96) involved cerebellar/subcortical, temporoparietal, basal-ganglia, visual-occipital, and insular white-matter territories.

### Amyloid component maps show stronger correspondence with rsfMRI-derived intrinsic network organization than Tau maps

Comparison with NeuroMark 2.2 intrinsic functional connectivity templates showed a clear modality difference in spatial correspondence across the 96 joint components, including both co-localized and modality-localized components. Fifty Amyloid component maps and 27 Tau component maps showed spatial correspondence above r = 0.40, whereas 46 Amyloid maps and 69 Tau maps were below this level. Overall, Amyloid component maps more frequently showed appreciable correspondence with rsfMRI-derived intrinsic network organization than Tau component maps.

### Joint component loadings show distinct disease-stage patterns across co-localized and modality-localized components

Linear mixed-effects models of subject-level joint ICA loadings showed diagnosis-related differences across multiple covarying Amyloid-Tau components after adjustment for age, sex, education, race, imaging site, and repeated observations. Figure 3 displays the signed significance of each diagnosis contrast, where the sign reflects the direction of the loading difference and the magnitude is shown on a -log10(q) scale.

**Figure 3.**
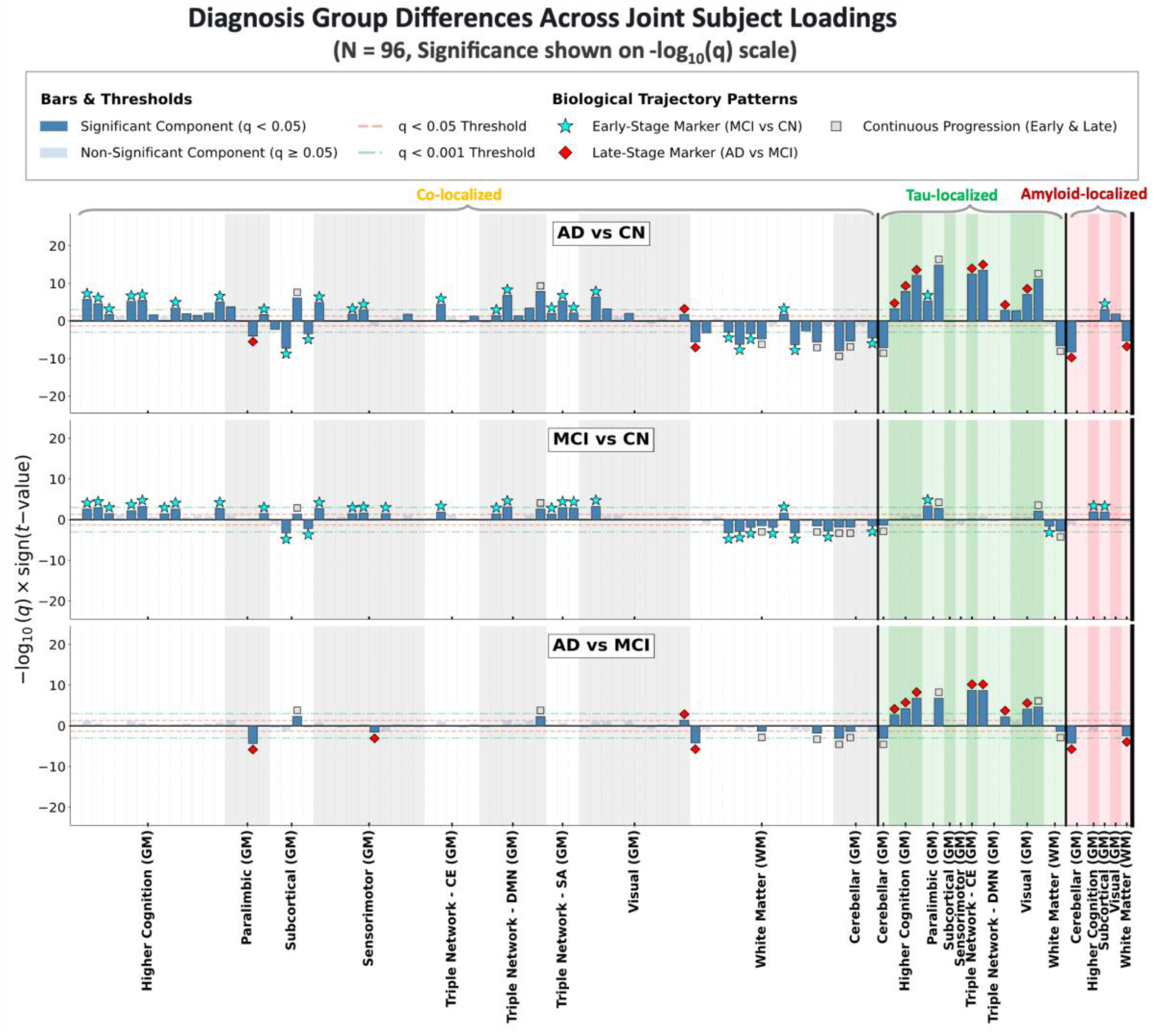
Diagnosis-stage differences in joint ICA loadings. Component-wise linear mixed-effects models tested MCI versus CN, AD versus CN, and AD versus MCI contrasts while adjusting for age, sex, education, race, imaging site, and repeated observations. Values are shown on a signed -log10(q) scale, where q denotes the FDR-corrected p-value and the sign indicates the direction of the group difference.

After FDR correction, 23 components showed significant loading differences for AD versus MCI, 44 for MCI versus CN, and 66 for AD versus CN. The direction of each difference was determined from the estimated diagnosis contrast in the mixed-effects model, with positive contrasts indicating higher joint loadings in the more advanced diagnostic group and negative contrasts indicating lower loadings (Figure 3). Among the significant components, positive loading differences were observed across several gray-matter systems, including higher-cognition, sensorimotor, visual, and triple-network domains, whereas negative differences were more prominent in a subset of subcortical, cerebellar, and white-matter components. Thus, diagnosis-related changes in joint component expression were spatially heterogeneous rather than reflecting a uniform increase or decrease across all components.

The distribution of significant components differed across disease-stage contrasts and across the spatial component groups. In the MCI-versus-CN comparison, significant differences were observed predominantly among co-localized components, spanning higher-cognition, subcortical, sensorimotor, triple-network, visual, and white-matter territories (Figure 3, middle panel). In contrast, the AD-versus-MCI comparison showed fewer significant co-localized components and a greater concentration of differences among Tau-localized (Figure 3, lower panel). The AD-versus-CN contrast showed the broadest separation, with 66 significant components distributed across both co-localized and modality-localized groups (Figure 3, upper panel).

### Pathology burden compartments revealed distinct disease-stage patterns within joint networks

When summarized across the 96 component territories, Amyloid-only, Tau-only, and dual-pathology burden showed distinct disease-stage profiles (Figure 4A). Amyloid-only burden increased from CN to MCI and showed comparatively little additional increase from MCI to AD. Tau-only burden showed a later-stage profile, with the larger increase occurring at the AD stage. Dual-pathology burden increased across the clinical spectrum and showed the largest overall CN- to-AD expansion. Averaged across components, the mean CN-to-AD increase was approximately 5.2 percentage points of component territory for Amyloid-only burden, 8.6 percentage points for Tau-only burden, and 13.7 percentage points for dual-pathology burden.

**Figure 4.**
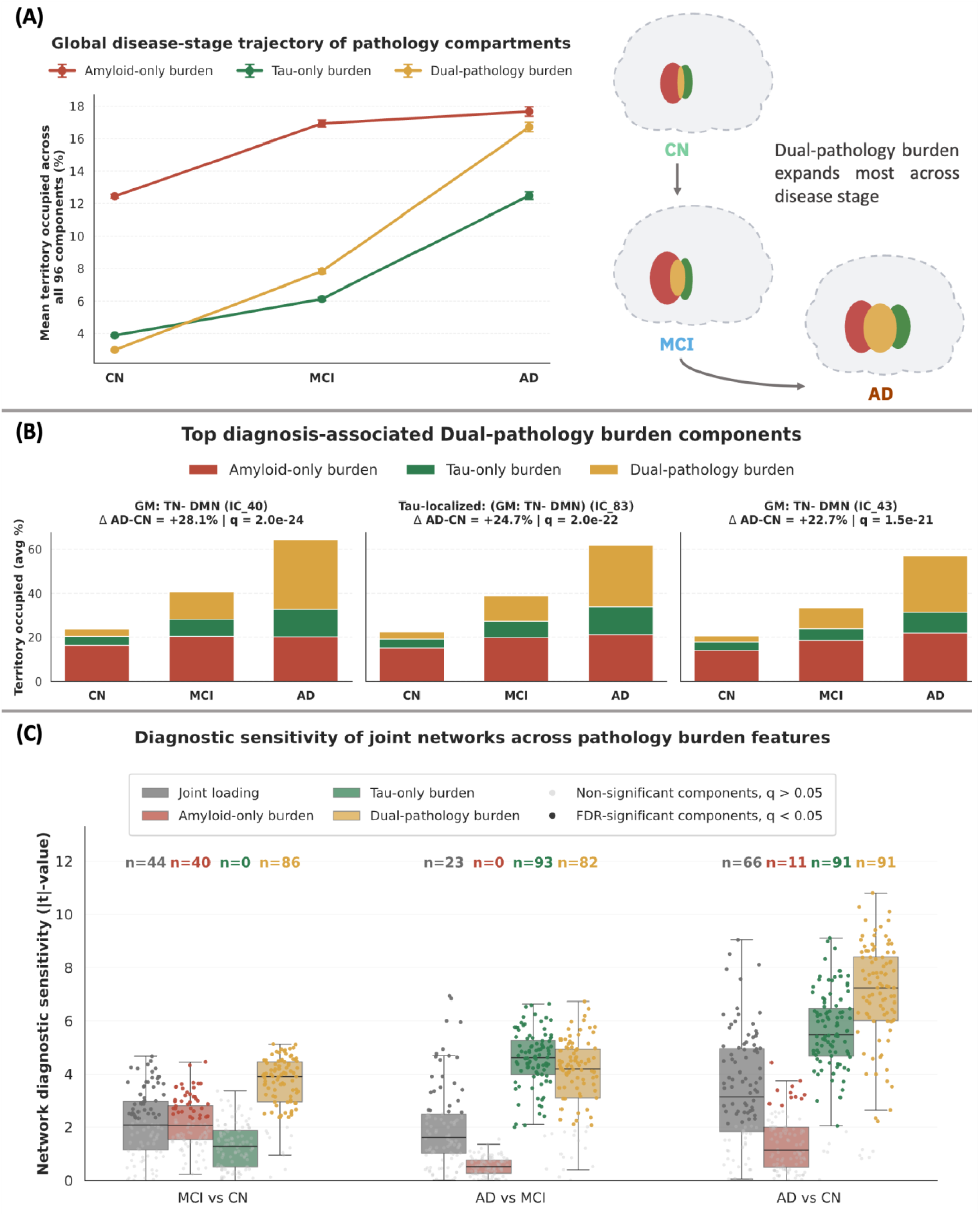
Spatially resolved pathology burden profiles across joint Amyloid-Tau component territories and diagnosis groups. (A) mean Amyloid-only, Tau-only, and dual-pathology burden across CN, MCI, and AD, averaged across the 96 joint component territories after first averaging subjects within each component and diagnosis group. The schematic illustrates the increasing overlap of Amyloid and Tau abnormality across diagnosis groups. (B) Representative components with the strongest AD-versus-CN dual-pathology burden increases among FDR-significant components. Stacked bars show the mean percentage of each component territory occupied by mutually exclusive Amyloid-only, Tau-only, and dual-pathology burden compartments. The three largest increases were observed in default-mode-related components. (C) Component-wise diagnosis differences for joint loading and the three pathology burden features. Boxplots show the distribution of absolute LMM t-values across components for each diagnosis contrast, with points representing individual components. Labels indicate the number of FDR-significant components for each feature and contrast.

The strongest CN-to-AD increases in dual-pathology burden were concentrated in default-mode components. Among components with FDR-significant AD-versus-CN effects, the three largest dual-pathology increases were IC40 (TN-DMN), IC83 (Tau-localized TN-DMN), and IC43 (TN-DMN) (Figure 4B). This localization provides spatial context for the global disease-stage pattern and identifies specific data-driven coupling territories in which overlapping Amyloid-Tau abnormality was particularly prominent.

Component-wise diagnosis contrasts further differentiated the three pathology burden compartments across the high-order joint component territories (Figure 4C). For MCI versus CN, FDR-significant differences were observed in 44 components for joint loading, 40 for Amyloid-only burden, none for Tau-only burden, and 86 for dual-pathology burden. For AD versus MCI, the corresponding numbers were 23, 0, 93, and 82 components, respectively; for AD versus CN, they were 66, 11, 91, and 91 components. Thus, the spatial distribution of diagnosis effects differed markedly across burden types: Amyloid-only differences were concentrated in the MCI-versus-CN contrast, whereas Tau-only differences were widespread in contrasts involving AD. Dual-pathology burden also showed broadly distributed diagnosis effects across disease stages, involving 86 components for MCI versus CN, 82 for AD versus MCI, and 91 for AD versus CN. Notably, it showed the broadest distribution in the MCI-versus-CN contrast, whereas Tau-only burden involved more components in AD versus MCI and an equal number in AD versus CN.

Relative to joint loading, the burden measures provided a more differentiated spatial profile of pathology across disease stages. Dual-pathology burden showed significant diagnosis effects in more component territories than joint loading for MCI versus CN (86 vs 44), AD versus MCI (82 vs 23), and AD versus CN (91 vs 66), while Amyloid-only and Tau-only burden showed distinct early- and later-stage distributions across the same component space. These component-wise patterns complement the average burden profiles in Figure 4A and the regional dual-pathology increases shown in Figure 4B.

### Dual pathology burden showed the strongest cognition association

ADAS13 associations differed across the four component-level features (Figure 5A). The number of FDR-significant components was highest for dual-pathology burden (86 components), followed by Tau-only burden (42 components) and joint loading (13 components). Amyloid-only burden showed no FDR-significant ADAS13 associations.

**Figure 5.**
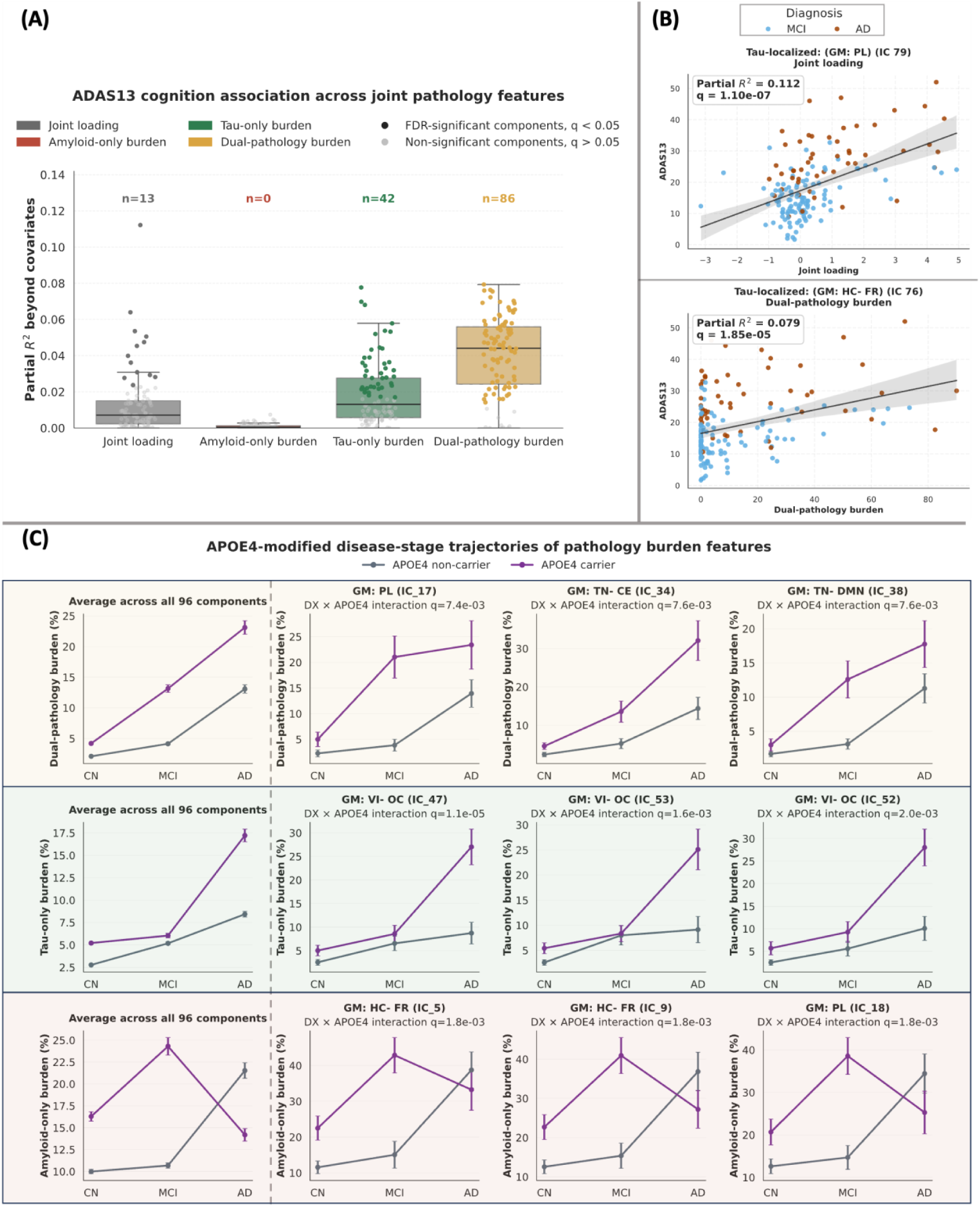
Cognition associations and APOE4-dependent pathology burden profiles. (A) Component-wise ADAS13 association strength for joint loading, Amyloid-only burden, Tau-only burden, and dual-pathology burden; partial R² represents feature-specific association strength beyond demographic, diagnosis, and imaging-site covariates. (B) Representative associations for the two component features with the highest ADAS13 partial R²; points may display MCI and AD for visualization, but reported statistics corresponds to the full adjusted analysis. (C) APOE4-stratified disease-stage profiles for dual-pathology, Tau-only, and Amyloid-only burden. The first column summarizes mean profiles across all 96 components, and the remaining columns show the three strongest diagnosis-by-APOE4 interactions for each burden feature. Points represent diagnosis-group means and error bars represent SEM. APOE4 analysis excludes participants with unavailable genotype.

The partial *R*^2^distribution showed the same pattern of association strength. Dual pathology burden had the highest median partial *R*^2^ values, indicating that it explained the greatest additional ADAS13 variance beyond age, sex, education, race, diagnosis, and imaging site covariates. Tau-only burden showed an intermediate association pattern, whereas joint loading had fewer significant associations and Amyloid-only burden remained near zero across components. Overall, cognitive impairment was most strongly related to the proportion of joint network territory occupied by simultaneous Amyloid and Tau abnormality.

### APOE4 is associated with different disease-stage profiles of pathology composition

APOE4 carrier status showed significant diagnosis-by-APOE4 interaction effects on pathology burden across multiple components (Figure 5C). Amyloid-only burden and dual-pathology burden showed comparable numbers of FDR-significant interaction effects, with 81 and 79 components, respectively, whereas Tau-only burden showed fewer interaction effects (18 components). This pattern indicates that APOE4-related disease-stage differences were more evident for Amyloid-containing burden features than for isolated Tau-only burden.

Figure 5C summarizes these interaction patterns across burden features. The first column shows the average APOE4-stratified disease-stage profile across all 96 components, while the remaining columns show the top three components for each burden feature, ranked by the FDR-corrected diagnosis-by-APOE4 interaction q-value. For dual-pathology burden, APOE4 carriers showed higher burden and steeper increases across disease stage than non-carriers, with the clearest separation emerging from MCI to AD. Tau-only burden showed fewer significant interactions overall, although the strongest components still showed APOE4-dependent differences across diagnosis groups. Amyloid-only burden showed a different pattern, with several components showing reduced or flattened disease-stage profiles among APOE4 carriers at later stages. Because Amyloid-only burden excludes voxels where Tau is also abnormal, this pattern may reflect a shift from isolated Amyloid abnormality toward overlapping Amyloid-Tau abnormality in APOE4 carriers at more advanced disease stages. Together, these results show that APOE4 carrier status was associated with different diagnosis-stage patterns of pathology burden, particularly for burden features involving Amyloid abnormality.

## Discussion

This study provides a fine-grained, data-driven map of cross-subject Amyloid-Tau PET coupling and separates two aspects of paired pathology that are often conflated: the spatial organization of covariance between the two tracers and the subject-specific composition of pathology within those spatial territories. High-order joint ICA served as the spatial discovery framework, identifying components in which Amyloid and Tau varied together across paired sessions. The resulting component territories were then used to quantify how much of each region was occupied by Amyloid-only, Tau-only, or simultaneous Amyloid-Tau abnormality. The central contribution is therefore not simply a larger ICA decomposition, but a spatially refined multimodal representation that moves from covariance discovery to pathology composition within the same data-driven units.

Prior Amyloid-Tau joint ICA work established that the two tracers can participate in linked but spatially distinct large-scale patterns (Pereira *et al*., 2019). The present findings extend that observation at a finer spatial scale. Rather than describing only a small set of broad covariance modes, the high-order solution resolved a larger set of localized component territories and showed that cross-subject Amyloid-Tau coupling can be expressed through both spatially corresponding and modality-localized patterns. This distinction refines the question from whether Amyloid and Tau are globally similar or different to how their statistical coupling is spatially organized across the brain. The high-order decomposition should therefore be viewed as a tool for spatial specificity, not as the biological endpoint of the study.

Mapping the PET-derived components to rsfMRI-derived intrinsic functional network templates provided a systems-level context for this spatial heterogeneity. Amyloid component maps more frequently showed strong correspondence with NeuroMark templates than Tau maps, and several of the strongest dual-pathology disease-stage effects occurred in default-mode territories. Prior work has linked early Amyloid accumulation to core default-mode regions and has shown that network architecture shapes regional Tau accumulation and propagation (Palmqvist *et al*., 2017; Franzmeier *et al*., 2020; Frontzkowski *et al*., 2022; Zheng *et al*., 2024). Our analysis does not demonstrate functional dysconnectivity directly because subject-specific fMRI connectivity was not modeled here. Rather, it suggests that data-driven Amyloid-Tau coupling territories are embedded within recognizable intrinsic brain systems and that the strength of this correspondence differs between the two tracer maps.

The burden analysis adds an interpretability layer to the joint decomposition. A joint loading measures how strongly a paired session expresses a particular multimodal covariance pattern, whereas pathology burden measures the within-territory spatial composition of abnormality in that session. These features are therefore complementary rather than competing biomarkers. The disease-stage profiles differed across the three mutually exclusive burden states: Amyloid-only burden was more evident in the earlier CN-to-MCI contrast, Tau-only burden in later-stage contrasts, and dual-pathology burden showed the broadest effects across the clinical spectrum. Because the burden compartments are mutually exclusive, an apparent plateau or reduction in Amyloid-only burden should not be interpreted as reduced Amyloid relevance; territory that becomes abnormal for both tracers is reassigned from Amyloid-only to dual-pathology burden.

Among the pathology compartments, dual-pathology burden showed the largest average CN-to-AD expansion, and the strongest effects were concentrated in default-mode components. This localization is consistent with the known vulnerability of default-mode and posterior association systems to AD pathology, but the present analysis adds a more specific observation: these systems emerged as data-driven Amyloid-Tau coupling territories in which spatially overlapping abnormality was particularly disease-relevant. The cognition analysis converged with this pattern, as dual-pathology burden showed the highest ADAS13 partial R² among the tested component features. Previous PET studies have shown that combined Amyloid and Tau positivity is associated with substantially greater risk of subsequent cognitive decline than Amyloid positivity alone (Ossenkoppele *et al*., 2022). The present results add spatial resolution to this broader observation by showing that, within the discovered coupling territories, cognitive association strength was greatest for the proportion of territory in which both tracer signals were abnormal.

The APOE4 analysis provided an additional test of whether pathology composition within the joint territories reflected known biological heterogeneity in AD. Diagnosis-by-APOE4 interactions were considerably more widespread for Amyloid-only and dual-pathology burden than for Tau-only burden, consistent with evidence that APOE4 potentiates the relationship between Amyloid and Tau pathology (Therriault *et al*., 2021). The APOE4 profiles should nevertheless be interpreted cautiously. Because Amyloid-only, Tau-only, and dual-pathology measures partition the same component territory, changes in one compartment necessarily affect the available territory for the others. The apparent late-stage reduction of Amyloid-only burden in some carrier profiles may therefore reflect redistribution toward dual-pathology burden rather than a decline in Amyloid involvement. Direct longitudinal testing of within-person compartment transitions is required before making a mechanistic claim.

A broader implication of this framework is that refined coupling territories provide candidate spatial units for future mechanistic modeling. Regional Amyloid-Tau interactions have been linked to subsequent Tau accumulation through both local and connectivity-mediated pathways (Lee *et al*., 2022), and recent connectomic models show that regional Amyloid burden improves the explanation of Tau deposition beyond connectivity alone (Zheng *et al*., 2024). The present decomposition cannot determine why Amyloid and Tau covary in a given territory, but it can identify where such covariance is expressed with greater spatial specificity. Longitudinal mediation or causal models could subsequently test whether shared network vulnerability, regional Amyloid burden, connectivity, or other biological factors precede and explain the emergence of Tau within specific coupling territories.

The results should be interpreted in light of several methodological considerations. First, model order is a scale parameter rather than a uniquely optimal biological solution. K = 100 was selected to obtain a high-order representation of spatial covariance, and ICA literature supports the use of higher orders to subdivide broad sources, but this evidence is derived largely from fMRI (Abou-Elseoud *et al*., 2010; Iraji *et al*., 2023). Model-order sensitivity should therefore be demonstrated empirically in the present PET data. Second, the classification of paired maps depends on Pearson spatial correlation. Spatial correlation captures topographic correspondence but is influenced by relative map intensity and background voxels; a thresholded overlap metric and permutation-based null would strengthen the co-localization interpretation. Third, burden estimates depend on the definition of component territory, the CN Aβ− reference group, and the voxelwise 95th-percentile threshold. Alternative spatial and abnormality thresholds may change absolute burden estimates. Finally, independent validation in another paired Amyloid-Tau PET cohort and direct comparison with whole-brain burden measures will be important for establishing the reproducibility and incremental spatial value of the component-level framework.

## Conclusion

By applying high-order joint ICA to paired Amyloid and Tau PET, this study establishes an unsupervised, data-driven framework for mapping fine-grained territories of cross-subject Amyloid-Tau coupling without imposing predefined spatial boundaries or forcing the two tracer maps to overlap. The resulting representation reveals that coupling is spatially heterogeneous and can be expressed through both shared and modality-localized patterns embedded within recognizable brain systems. Using these same data-derived territories to quantify pathology composition further shows how isolated and overlapping abnormalities can be described within a common spatial framework. This refined mapping approach provides a foundation for future longitudinal and mechanistic analyses aimed at understanding why Amyloid and Tau converge in particular brain territories and how that convergence relates to clinical progression.

## Data Availability Statement

Data used in this study were obtained from the Alzheimer’s Disease Neuroimaging Initiative (ADNI) database. ADNI imaging, clinical, demographic, and genetic data are available to qualified investigators through the ADNI data access portal hosted by the Laboratory of Neuro Imaging Image and Data Archive (LONI IDA), subject to ADNI data-use approval and policies. The datasets analyzed in this study can be accessed at the ADNI repository: adni.loni.usc.edu. Derived component maps, burden measures, and analysis outputs generated for this study are available from the corresponding author upon request, subject to applicable ADNI data-sharing restrictions.

## References

Abou-Elseoud, A. et al. (2010) “The effect of model order selection in group PICA,” Human Brain Mapping, 31(8), pp. 1207–1216. Available at: 10.1002/hbm.20929.

Benjamini, Y. and Hochberg, Y. (1995) “Controlling the False Discovery Rate: A Practical and Powerful Approach to Multiple Testing,” Journal of the Royal Statistical Society: Series B (Methodological), 57(1), pp. 289–300. Available at: 10.1111/j.2517-6161.1995.tb02031.x.

Bimali, B. et al. (2025) “Multimodal Fusion Analysis of [18F]Florbetapir PET and Multiscale Functional Network Connectivity in Alzheimer’s Disease,” 2025 IEEE EMBS International Conference on Biomedical and Health Informatics (BHI). 2025 IEEE EMBS International Conference on Biomedical and Health Informatics (BHI), pp. 1–6. Available at: 10.1109/BHI67747.2025.11269510.

Calhoun, V. d., et al. (2006) “Method for multimodal analysis of independent source differences in schizophrenia: Combining gray matter structural and auditory oddball functional data,” Human Brain Mapping, 27(1), pp. 47–62. Available at: 10.1002/hbm.20166.

Chen, J. et al. (2026) “Dynamic Fusion of Genomics and Functional Network Connectivity in UK Biobank Reveals Schizophrenia-Related SNP Manifolds,” Human Brain Mapping, 47(6), p. e70530. Available at: 10.1002/hbm.70530.

Cho, H. et al. (2016) “In vivo cortical spreading pattern of tau and amyloid in the Alzheimer disease spectrum,” Annals of Neurology, 80(2), pp. 247–258. Available at: 10.1002/ana.24711.

Clark, C.M. et al. (2011) “Use of florbetapir-PET for imaging beta-amyloid pathology,” JAMA, 305(3), pp. 275–283. Available at: 10.1001/jama.2010.2008.

Fonseca, C.S. et al. (2026) “Spatial patterns of tau accumulation across the Alzheimer’s disease spectrum,” Alzheimer’s & Dementia: The Journal of the Alzheimer’s Association, 22(1), p. e71076. Available at: 10.1002/alz.71076.

Franzmeier, N. et al. (2020) “Patient-centered connectivity-based prediction of tau pathology spread in Alzheimer’s disease,” Science Advances, 6(48), p. eabd1327. Available at: 10.1126/sciadv.abd1327.

Frontzkowski, L. et al. (2022) “Earlier Alzheimer’s disease onset is associated with tau pathology in brain hub regions and facilitated tau spreading,” Nature Communications, 13(1), p. 4899. Available at: 10.1038/s41467-022-32592-7.

Himberg, J. and Hyvarinen, A. (2003) “Icasso: software for investigating the reliability of ICA estimates by clustering and visualization.” 2003 IEEE XIII Workshop on Neural Networks for Signal Processing (IEEE Cat. No.03TH8718), IEEE, p. 27. Available at: 10.1109/NNSP.2003.1318025.

Iraji, A. et al. (2023) “Identifying canonical and replicable multi-scale intrinsic connectivity networks in 100k+ resting-state fMRI datasets,” Human Brain Mapping, 44(17), pp. 5729–5748. Available at: 10.1002/hbm.26472.

Jack, C.R. et al. (2018) “NIA-AA Research Framework: Toward a biological definition of Alzheimer’s disease,” Alzheimer’s & Dementia: The Journal of the Alzheimer’s Association, 14(4), pp. 535–562. Available at: 10.1016/j.jalz.2018.02.018.

Jack Jr., C.R., et al. (2024) “Revised criteria for diagnosis and staging of Alzheimer’s disease: Alzheimer’s Association Workgroup,” Alzheimer’s & Dementia, 20(8), pp. 5143–5169. Available at: 10.1002/alz.13859.

Jensen, K.M. et al. (2024) “Addressing Inconsistency in Functional Neuroimaging: A Replicable Data-Driven Multi-Scale Functional Atlas for Canonical Brain Networks,” bioRxiv, p. 2024.09.09.612129. Available at: 10.1101/2024.09.09.612129.

Johnson, K.A. et al. (2016) “Tau positron emission tomographic imaging in aging and early Alzheimer disease,” Annals of Neurology, 79(1), pp. 110–119. Available at: 10.1002/ana.24546.

Landau, S.M. et al. (2012) “Amyloid deposition, hypometabolism, and longitudinal cognitive decline,” Annals of Neurology, 72(4), pp. 578–586. Available at: 10.1002/ana.23650.

Landau, S.M. et al. (2024) “Positron emission tomography harmonization in the Alzheimer’s Disease Neuroimaging Initiative: A scalable and rigorous approach to multisite amyloid and tau quantification,” Alzheimer’s & Dementia, 21(1), p. e14378. Available at: 10.1002/alz.14378.

Lee, W.J. et al. (2022) “Regional Aβ-tau interactions promote onset and acceleration of Alzheimer’s disease tau spreading,” Neuron, 110(12), pp. 1932–1943.e5. Available at: 10.1016/j.neuron.2022.03.034.

Ossenkoppele, R. et al. (2022) “Amyloid and tau PET-positive cognitively unimpaired individuals are at high risk for future cognitive decline,” Nature Medicine, 28(11), pp. 2381–2387. Available at: 10.1038/s41591-022-02049-x.

Ossenkoppele, R., van der Kant, R. and Hansson, O. (2022) “Tau biomarkers in Alzheimer’s disease: towards implementation in clinical practice and trials,” The Lancet. Neurology, 21(8), pp. 726–734. Available at: 10.1016/S1474-4422(22)00168-5.

Palmqvist, S. et al. (2017) “Earliest accumulation of β-amyloid occurs within the default-mode network and concurrently affects brain connectivity,” Nature Communications, 8(1), p. 1214. Available at: 10.1038/s41467-017-01150-x.

Pereira, J.B., et al. (2019) “Amyloid and tau accumulate across distinct spatial networks and are differentially associated with brain connectivity,” eLife. Edited by M. Irish and T.E. Behrens, 8, p. e50830. Available at: 10.7554/eLife.50830.

Pontecorvo, M.J. et al. (2019) “A multicentre longitudinal study of flortaucipir (18F) in normal ageing, mild cognitive impairment and Alzheimer’s disease dementia,” Brain: A Journal of Neurology, 142(6), pp. 1723–1735. Available at: 10.1093/brain/awz090.

St-Onge, F. et al. (2024) “Tau accumulation and its spatial progression across the Alzheimer’s disease spectrum,” Brain Communications, 6(1), p. fcae031. Available at: 10.1093/braincomms/fcae031.

Sui, J. et al. (2012) “A Review of Multivariate Methods for Multimodal Fusion of Brain Imaging Data,” Journal of neuroscience methods, 204(1), pp. 68–81. Available at: 10.1016/j.jneumeth.2011.10.031.

Therriault, J. et al. (2021) “APOEε4 potentiates the relationship between amyloid-β and tau pathologies,” Molecular Psychiatry, 26(10), pp. 5977–5988. Available at: 10.1038/s41380-020-0688-6.

Vogel, J.W. et al. (2020) “Spread of pathological tau proteins through communicating neurons in human Alzheimer’s disease,” Nature Communications, 11(1), p. 2612. Available at: 10.1038/s41467-020-15701-2.

Zheng, L. et al. (2024) “Combined Connectomics, MAPT Gene Expression, and Amyloid Deposition to Explain Regional Tau Deposition in Alzheimer Disease,” Annals of Neurology, 95(2), pp. 274–287. Available at: 10.1002/ana.26818.

